# Insect herbivory on restored rainforest seedlings weakened by neighbours but unaffected by invasive coffee

**DOI:** 10.1101/2025.07.18.665266

**Authors:** Lakshmi Niranjana, Aparna Krishnan, Akhil Murali, T. R. Shankar Raman, Divya Mudappa, Mahesh Sankaran, Mayank Kohli

**Affiliations:** National Centre for Biological Sciences, Tata Institute of Fundamental Research, Bellary Road, Bengaluru 560065, Karnataka, India; Nature Conservation Foundation, 1311, 12th A Main, Vijayanagar 1st Stage, Mysuru, 570017, Karnataka, India; Institute of Ecology, Leuphana University of Lüneburg, Lüneburg, Germany; Institute of Biology/Geobotany and Botanical Garden, Martin Luther University, Halle-Wittenberg, Halle (Saale), Germany

**Keywords:** Associational effects, coffee invasion, plant traits, tropical forest restoration, Western Ghats

## Abstract

Restoration of degraded tropical forests is often impeded by invasive species. For regenerating native seedlings, the presence of invasives in the neighbourhood can alter insect herbivory patterns, ultimately shaping restoration trajectories; however, such indirect effects are rarely examined. Here, we investigated the effect of robusta coffee (*Coffea canephora*) - a shade-tolerant invasive species under closed-canopy secondary forests in the Western Ghats – on the incidence of herbivory (proportion of leaves with any sign of damage) and the extent of leaf damage (percentage leaf area consumed) in seedlings of 10 rainforest species in plots from which coffee plants were either weeded out or left intact. We further examined whether local neighbourhood densities of coffee and other saplings, and species’ leaf traits, explained herbivory patterns. Removal of invasive coffee did not influence herbivory incidence or the extent of damage across our focal species. However, the incidence of herbivory declined with increasing neighbourhood plant density, suggesting that neighbourhood plants provide a resource dilution effect. Both incidence and extent of herbivore damage were strongly species-specific and partly explained by leaf traits: greater leaf carbon content was correlated with lower herbivory incidence. Contrary to expectations, plants with resource-acquisitive traits (high leaf nitrogen and specific leaf area) experienced lower incidence of herbivory and extent of damage. Our findings suggest that in this system, the indirect effects via herbivores are perhaps not as important in influencing restoration as the more direct effects of invasive species, such as competition for resources.

## 1. INTRODUCTION

Invasive plant species are among the greatest threats to the recovery of degraded tropical forests (Dawson, Burslem, & Hulme, 2015; Döbert, Webber, Sugau, Dickinson, & Didham, 2018; Waddell et al., 2020, 2023; Weidlich, Flórido, Sorrini, & Brancalion, 2020). Manual removal of invasive plants, an important restoration strategy in site preparation for native regeneration, is intensive in effort, time, and cost (Kettenring & Adams, 2011). Understanding whether and how invasives limit the recovery of native species and experiments that assess the effects of different removal techniques on native regeneration can help create suitable restoration plans (Flory & Clay, 2009) and arrive at cost-effective decisions on weed removal during forest restoration. Presently, little is known about whether shade-tolerant invasive species that persist under relatively closed canopies limit native regeneration in degraded and secondary forests and the mechanisms by which they do so (Martin, Canham, & Marks, 2009). This gap is particularly concerning as evidence from multiple tropical regions demonstrates that secondary forests often reach a state of apparent arrested succession of only partial recovery toward benchmark forest structure and composition (Arroyo-Rodríguez et al., 2017; Goldsmith et al., 2011), with shade-tolerant invasives continuing to increase in abundance through succession (Joshi, Mudappa, & Raman, 2009; Martin et al., 2009). Understanding the mechanisms through which shade-tolerant invasives limit regeneration and whether such forests require continuous invasive-species management can inform forest restoration efforts.

Arrested succession due to persistent invasives can result from both direct and indirect mechanisms. Direct mechanisms include resource competition, as seen in *Acer platanoides*, which reduces photosynthetically active radiation (PAR) in the understory by up to 55%, significantly lowering native plant survival in invaded riparian zones (Reinhart, Gurnee, Tirado, & Callaway, 2006). Similarly, invasive plants can also directly alter microclimatic conditions such as temperature and light availability to modify native regeneration (Garcia & Clusella-Trullas, 2025). Indirect effects of invasives on forest tree communities include top-down control of vegetation by herbivores and pathogens. These effects are particularly pronounced in tropical regions, where herbivory rates are considerably higher compared to temperate forests, with most leaf damage occurring during early development when young and expanding leaves are most vulnerable (Aide, 1993; P. D. Coley & Barone, 1996). In tropical rainforests, top-down control by plant consumers such as insect herbivores and fungal pathogens plays a key role in determining seedling establishment and the composition of tree communities (Bagchi et al., 2014; Krishnadas, Bagchi, Sridhara, & Comita, 2018). Invasive plants can indirectly modify such top-down effects by herbivory and pathogens through associational effects wherein the diversity and composition of surrounding vegetation are thought to modify the interaction between focal plants and herbivores (Barbosa et al., 2009a; Hambäck, Inouye, Andersson, & Underwood, 2014a; Klapwijk & Bonsall, 2022; Mutz et al., 2022a). When the palatability of an invasive species differs from that of neighbouring native species, four distinct scenarios can result (Hahn & Orrock, 2016). If the invasive species is unpalatable, it may either shield native plants from herbivory (associational resistance) or inadvertently concentrate herbivore feeding on the remaining palatable natives (associational selection). Conversely, if an invasive species is more palatable than a native species, it may either enhance herbivory pressure on natives by attracting herbivores (associational susceptibility) or reduce damage to natives by diverting herbivores toward themselves (associational escape). While relative palatability is a critical driver, total density of neighbouring plants can also alter herbivory patterns by either increasing herbivory pressure (resource concentration effects; Root, 1973) or by reducing its effects (resource dilution effects; Otway, Hector, & Lawton, 2005). All these outcomes can be mediated by local context (Kim, 2017), such as the density of neighbours or physical and chemical traits of invasives, which can intensify or buffer these effects. For instance, a considerable body of evidence shows that neighbourhood composition affects insect herbivory on focal plants (Jia et al., 2023; Kim, 2017; Stastny & Agrawal, 2014), although studies have also found inconsistent patterns (Zhang, Xu, Zhang, Zhang, & Cao, 2023), suggesting the context-specific influence of plant traits, herbivore preferences, and other environmental conditions.

Plant susceptibility or resistance to herbivory can also vary with chemical traits such as leaf chemical (e.g., nitrogen, carbon concentrations) and morphological traits (e.g., leaf dry matter content (LDMC), specific leaf area (SLA), leaf thickness; Caldwell, Read, & Sanson, 2016; Coley & Barone, 1996; Poorter, van de Plassche, Willems, & Boot, 2004a; Schuldt et al., 2012). Plant traits can thus help understand how herbivore-mediated indirect effects of invasive plant species on native plants are likely to vary across species. For example, herbivorous insects may preferentially consume palatable, nutritious species with large leaves (Carmona, Lajeunesse, & Johnson, 2011; Muiruri et al., 2019), or early successional species with high nutrient contents and low investment in defence compounds as opposed to late successional species with greater investment in defensive compounds (Poorter, van de Plassche, Willems, & Boot, 2004b; Schowalter, 1981; Zhao & Chen, 2012). Despite a rising number of studies that have focused on plant functional traits and the impacts of neighbourhood effects (Felix, Stevenson, & Koricheva, 2023; Mutz et al., 2022b; Wang et al., 2024), it is still unclear how differences in plant species diversity and density, and different chemical and morphological traits influence the patterns of insect herbivory.

In this study, we investigated the effects of coffee invasion on insect herbivory in tropical tree seedlings within a rainforest restoration experiment in the Western Ghats, India, a global biodiversity hotspot. Invasion by coffee is a major impediment to forest restoration and regeneration in the tropical rainforests of the region. Both mature and secondary rainforest patches in the region have been invaded by robusta coffee (*Coffea canephora*) and, to a lesser extent, by arabica coffee (*C. arabica*), displacing natural vegetation (Joshi, Mudappa, & Raman, 2009). Prior research suggests that coffee is a strong competitor in secondary forests as coffee plants have persisted in abandoned plantations even after more than 70 years (Raymundo et al., 2018). Given coffee’s distinct chemical traits, shade-tolerant characteristics (Rodríguez-López et al., 2014), and the relatively high density of saplings in invaded areas (Joshi, Mudappa, & Raman, 2009), its presence can also potentially influence herbivore pressure on native plants, in turn influencing vegetation community dynamics. The presence of coffee may influence insect herbivory patterns by either attracting or repelling herbivores, depending on the palatability of coffee plants relative to the native vegetation. However, while the direct effects of coffee invasion on native plant communities in tropical rainforests have been described (Joshi et al., 2009c), the potential indirect effects of such invasions are less well understood.

This research aimed to answer two specific questions: a) How does the presence and removal of invasive coffee affect insect herbivory on native tropical tree seedlings planted for restoration? and b) Do neighbourhood plant density and leaf traits of directly seeded native tree saplings predict their susceptibility to insect herbivory? To determine whether the presence of coffee influences insect herbivory on native plants through associational effects, we compared herbivory patterns on seedlings of ten directly-seeded native tree species in patches where invading coffee plants were manually removed with those in patches where coffee plants were left intact.

## 2. METHODS

### 2.1 Study Site

Our study was conducted in March 2024 on the Valparai Plateau, Tamil Nadu, India (10.26°– 10.37° N and 76.87°–76.99° E), situated in the Anamalai Hills of the Western Ghats and the Sri Lanka global biodiversity hotspot (Figure 1a). The plateau spans an area of 220 km² and is characterised by a mosaic of commercial plantations, including coffee, tea and cardamom, interspersed with fragments of rainforest and surrounded by four protected areas. It is a key biodiversity conservation site in the southern Western Ghats. The elevation of the Valparai Plateau ranges from 800 – 1500 m above mean sea level. The natural vegetation of the plateau is classified as mid-elevation tropical wet evergreen forest, specifically of the *Cullenia exarillata* – *Mesua ferrea* – *Palaquium ellipticum* type (Pascal 1988). The Valparai Plateau, once covered in tropical rainforest, was extensively cleared between 1890 and 1920 to establish tea and coffee plantations, which now dominate the landscape. Around 1,000 ha of rainforest persist on the Valparai Plateau, scattered across 80 fragments ranging from 0.5 ha to 300 ha. Many of these fragments have undergone degradation due to past deforestation, fragmentation, and conversion into plantations, some of which were later abandoned. Detailed information on the study site and restoration efforts can be found in Osuri, Kasinathan, Raman & Mudappa (2024) and Mudappa & Raman (2007).

**Figure 1.**
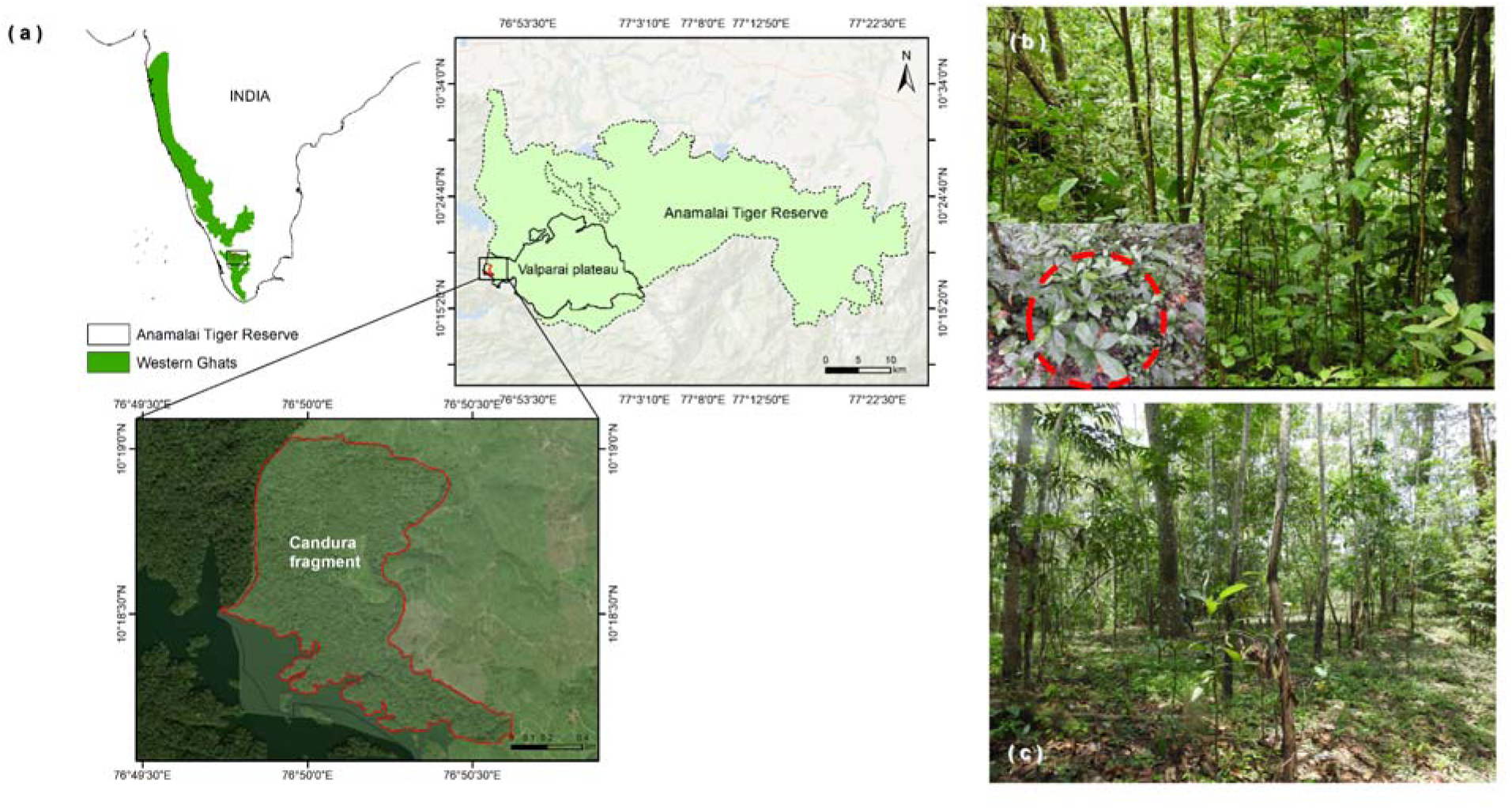
(a) Map of the Candura fragment within the Valparai Plateau in the Anamalai Tiger Reserve, Western Ghats, India. (b) Restoration plot in Candura invaded by coffee, showing a dense understory of coffee plants. (c) Coffee removal plot from the same restoration experiment, where invasive coffee plants have been cleared.

The restoration experimental site was a 7 ha area of the Candura rainforest fragment (10°18′31″ N, 76°50′02″ E; 875 m asl) on the western edge of the Valparai Plateau. The Candura fragment, shaped by episodic selective logging (most recently in the 1990s and early 2000s, and last recorded in 2004), includes abandoned coffee plantations where coffee (primarily robusta, *C. canephora*) has invaded secondary forests. Initially set aside as a smaller restoration site in 2007, Candura was later expanded to 121 ha for conservation and restoration efforts, with the area divided into 1 ha (100 × 100 m) grid cells for systematic restoration, research, and monitoring.

### 2.2 Experimental setup

The study utilised a rainforest restoration experiment to compare herbivory patterns under two different restoration treatments in sites invaded by robusta coffee: (1) direct seeding of native species without coffee removal (Figure 1b), and (2) direct seeding of native rainforest species after removal of invasive coffee (Figure 1c). A total of 22 plots (20 × 20 m) were established within 1-hectare grid cells in areas of similar coffee density and canopy openness. These plots were randomly assigned to the two treatments, with 11 replicates for each (Figure S1). Plots were weeded in October 2021, and direct seeding was carried out starting January 2022 using seeds of ten native species: *Canarium strictum*, *Cullenia exarillata*, *Heynea trijuga*, *Mesua ferrea*, *Litsea nigrescens*, *Cinnamomum malabatrum*, *Actinodaphne wightiana*, *Myristica beddomei*, *Palaquium ellipticum*, and *Machilus glaucescens* at a density of 1 seed / m2. Seeding was completed for nine species within 2022; for *Myristica beddomei*, however, seeding was conducted in both 2022 and 2023 due to limited seed availability.

Insect herbivory on leaves was assessed by quantifying the percentage of leaf area lost to insects. For this study, five individual seedlings of each species were haphazardly selected from each replicate plot to quantify herbivory. As most individuals were young, their height was <1 m at the time of measurement. For each focal species, we aimed to sample 5 leaves from 5 individual plants per plot (25 leaves per species per plot). However, due to constraints such as limited leaf availability in certain plots, sampling effort varied (15–25 leaves per species per plot). For species with compound leaves (*C. strictum*, *H. trijuga*), five random leaflets were selected from each plant. Photographs of the leaves were taken *in situ* using a Samsung S21 FE camera against a white portable background. These images were then analysed using the BioLeaf - Foliar Analysis™ mobile application, a non-destructive imaging method (Machado et al., 2016) to estimate the extent of the leaf area lost to insect herbivory. For all the leaves where the leaf contours were lost, we utilised interactive control points, creating a smooth contour that fits the original edge of the leaves to estimate the leaf area lost. Only mature, fully expanded leaves were sampled, while senescent and diseased leaves were excluded. The quantification of herbivory damage focused specifically on leaf mining, leaf chewing, and skeletonisation. Leaves exhibiting damage from mammalian herbivory or leaf curling were excluded from the study. For each focal individual in the herbivory estimation, neighbourhood plant density was quantified within a 1 m radius. Coffee plants and other neighbourhood plants (naturally regenerated) in the surrounding area were counted and categorised separately for analysis.

Leaf nitrogen, carbon, C: N ratio, leaf dry matter content (LDMC), specific leaf area (SLA) and leaf thickness were also quantified for the 10 focal species. Fully expanded and undamaged sun-exposed leaves from mature plants in the vicinity were selected for the quantification of traits. To prevent dehydration-induced shrinkage, samples were covered in wet paper and sealed in plastic bags post-collection. Within 24 hours, leaves were gently detached, patted dry, and scanned for area measurement analysis. The BlackSpot leaf area calculator software (Varma & Osuri, 2013) was used to estimate leaf area from leaf scans. Leaves were then oven-dried at 40°C for 72 hours to constant mass and weighed using a balance (precision: 0.001 g). SLA was calculated as leaf area (cm²) divided by dry mass (g), and LDMC was calculated as the ratio of leaf dry mass to fresh mass, expressed as mg/g. Leaf carbon and nitrogen percentages were quantified using a LECO Truspec CN elemental analyzer (LECO Corporation) on oven-dried, fine-powdered leaf samples.

### 2.3 Data analysis

Our raw data on herbivory damage were highly right-skewed and zero-inflated due to a large proportion of undamaged leaves. To account for this, we used a two-part hurdle model approach to separately analyse (i) the probability of herbivory incidence (i.e., whether or not a leaf showed any damage) and (ii) the extent of herbivory damage per affected individual.

We first modelled herbivory incidence as a binary response variable, where leaves were classified as either damaged (any leaf area damage) or undamaged (no leaf area missing). We fitted a generalised linear mixed-effects model (GLMM) with a binomial error distribution to test for differences in herbivory incidence between treatments (coffee presence vs coffee removal) across different species. The model included species and treatment (coffee removal vs presence) as fixed effects, with plot and individual as nested random effects. To further assess the effect of neighbourhood density on herbivory incidence, we fitted two additional GLMMs. The first model included the number of neighbouring coffee plants as a fixed effect, while the second model included the total number of neighbouring plants (both coffee and other species). In both models, we included an interaction term with species identity and retained the same nested random effects structure (plot and individual).

To test the difference in leaf area loss among attacked leaves between the two treatments (hereafter, extent of herbivory damage), we first removed all observations with zero leaf damage and modelled the proportion of leaf area lost only for leaves with damage present. Damage values were averaged across leaves at the individual plant level. Since the data were again rightly skewed (Figure S2), we log-transformed the response variable and fitted a linear mixed-effects model (LMM) with treatment, species, and their interaction as fixed effects and plot as a random effect to account for spatial autocorrelation. To assess whether neighbourhood density influences the extent of herbivory damage, we extended our analysis with two additional models. One accounted for the number of neighbouring coffee plants, and the other for the total number of neighbouring individuals (including both coffee and non-coffee species). In both cases, we tested for species-specific responses by including an interaction between the fixed effects, species identity and neighbourhood density, and included plot as a random effect to control for spatial structure.

To test whether leaf traits influenced susceptibility to herbivory, we focused on six functional traits measured for the ten focal species. Prior to analysis, we first examined pairwise correlations among traits to assess multicollinearity. Then we performed a principal component analysis (PCA) to reduce dimensionality and capture major axes of trait variation. The first two principal components, which together explained approximately 80% of the total variance, were retained as composite trait axes. These components were used as fixed effects in two separate models: (i) a generalised linear mixed model (GLMM) for herbivory incidence, which included plot and individual as nested random effects; and (ii) a linear mixed-effects model (LMM) for extent of herbivory damage, which included plot as a random effect.

We checked model assumptions using residual diagnostics (Q-Q plots, residuals vs fitted values). Predictor significance was tested using Type III Wald χ² tests (*car* package; Fox & Weisberg, 2019). All analyses used R v4.3.1 (R Core Team, 2023) with *glmer* for GLMMs and *lmer* for linear mixed-effect models from the lme4 package in R (Bates et al., 2015). Detailed model specifications are included in the supplementary tables.

## 3. RESULTS

### 3.1 General patterns of leaf herbivory

We sampled 3,953 leaves from 904 seedlings, of which 2545 leaves showed incidence of herbivory (N = 1311 in plots with coffee present; N = 1234 in plots where coffee was removed) across the 10 species. The extent of tissue loss to herbivores per seedling ranged from 0% to 42.7% of the leaf area, with 824 out of 904 seedlings showing some incidence of herbivory (N= 428 in coffee-present plots; N= 396 in coffee-removed plots). Average total neighbourhood density was 3.88 (±0.17) in plots with coffee present and 2.89 (±0.11) in plots where coffee had been removed. On average, neighbourhoods consisted of 1.33 (±0.14) coffee plants and 2.55 (±0.11) plants of other species in plots with coffee, and 0.21 (±0.04) coffee plants and 2.68 (±0.11) plants of other species in plots where coffee had been removed (Figure S3).

### 3.2 Effect of coffee removal and neighbourhood density on herbivory incidence and the extent of herbivory damage

Herbivory incidence did not differ significantly between treatments (Figure 2a; Table S1). Among seedlings that experienced herbivory, the extent of herbivory damage was slightly higher in plots from which coffee was removed (9.5%; 95% CI: 6.7–13.4%) compared to those where coffee was present (7.9%; 95% CI: 5.6–11.1%), but this difference was not statistically significant (Figure 2b; Table S2). There was no significant interaction between treatment and species identity for both herbivory incidence and extent of herbivory damage, indicating that species responses did not differ between plots with coffee present or removed.

**Figure 2.**
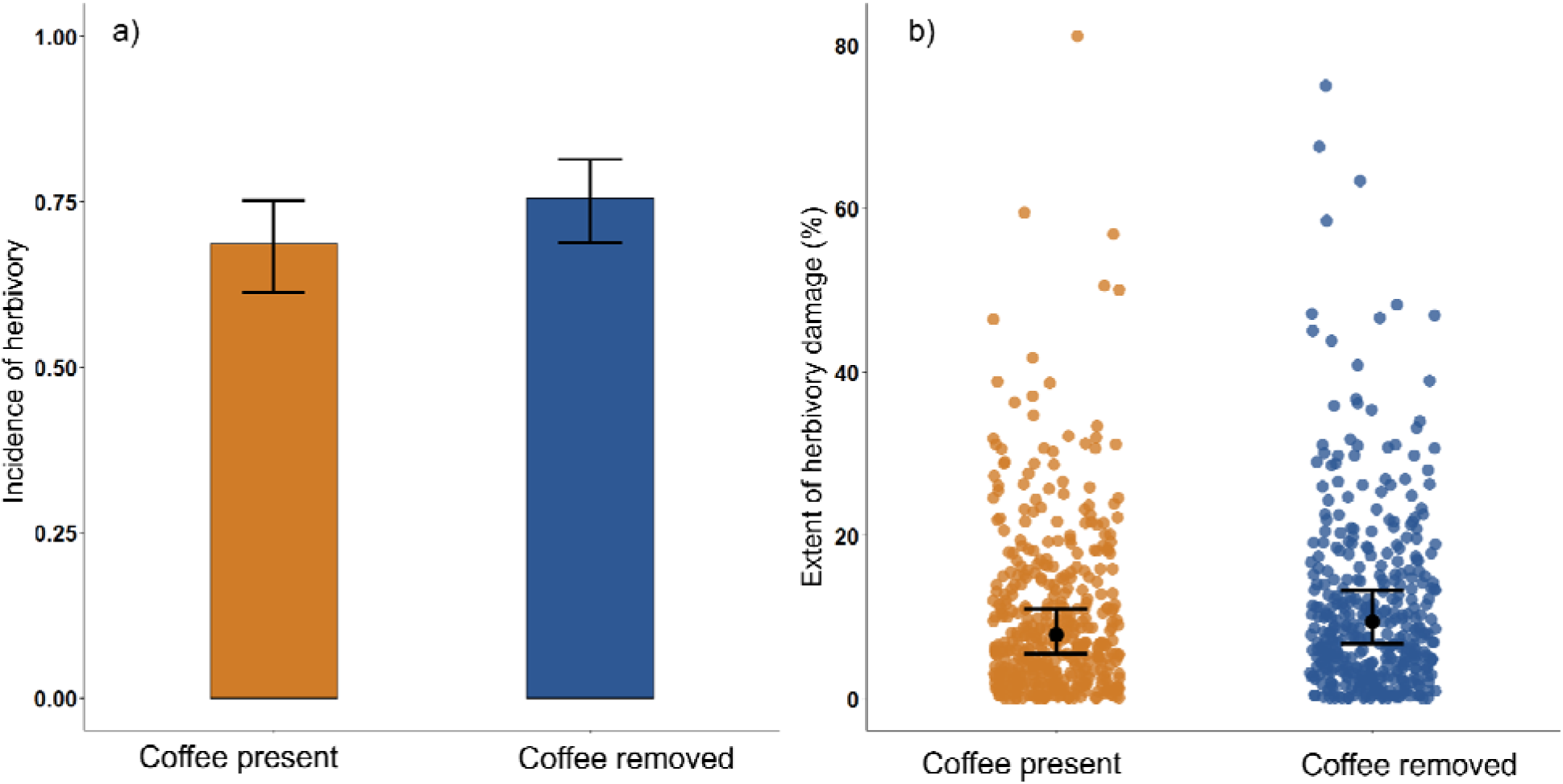
Predicted herbivory responses across treatments: (a) predicted incidence of herbivory (probability of any leaf damage) across coffee present (orange) and coffee removed (blue) treatments, and (b) observed (orange and blue dots) and predicted extent of herbivory damage (% leaf area lost); the black dots represent model-predicted means and the vertical bars (black) represent 95% confidence intervals. Predictions for the incidence of herbivory (a) were based on a generalised linear mixed-effects model (GLMM), while those for the extent of herbivory damage (b) were based on a linear mixed-effects model (LMM).

Although coffee removal did not affect herbivory, we found that the total density of plants in the 1m radius neighbourhood had a dilution effect on herbivory. The incidence of herbivory decreased as the total neighbourhood density (including both coffee and non-coffee species) increased, with incidence declining from 84.3 % to 80.3 % as neighbourhood densities increased from 0 to 5 neighbours (Figure 3; Table S3). Neighbourhood densities, however, did not affect the extent of herbivory damage (Table S4). Further, the density of neighbouring coffee plants, by themselves, did not affect either incidence or extent of herbivory damage, consistent with the findings that the coffee removal treatment did not impact herbivory (Tables S5 & S6).

**Figure 3.**
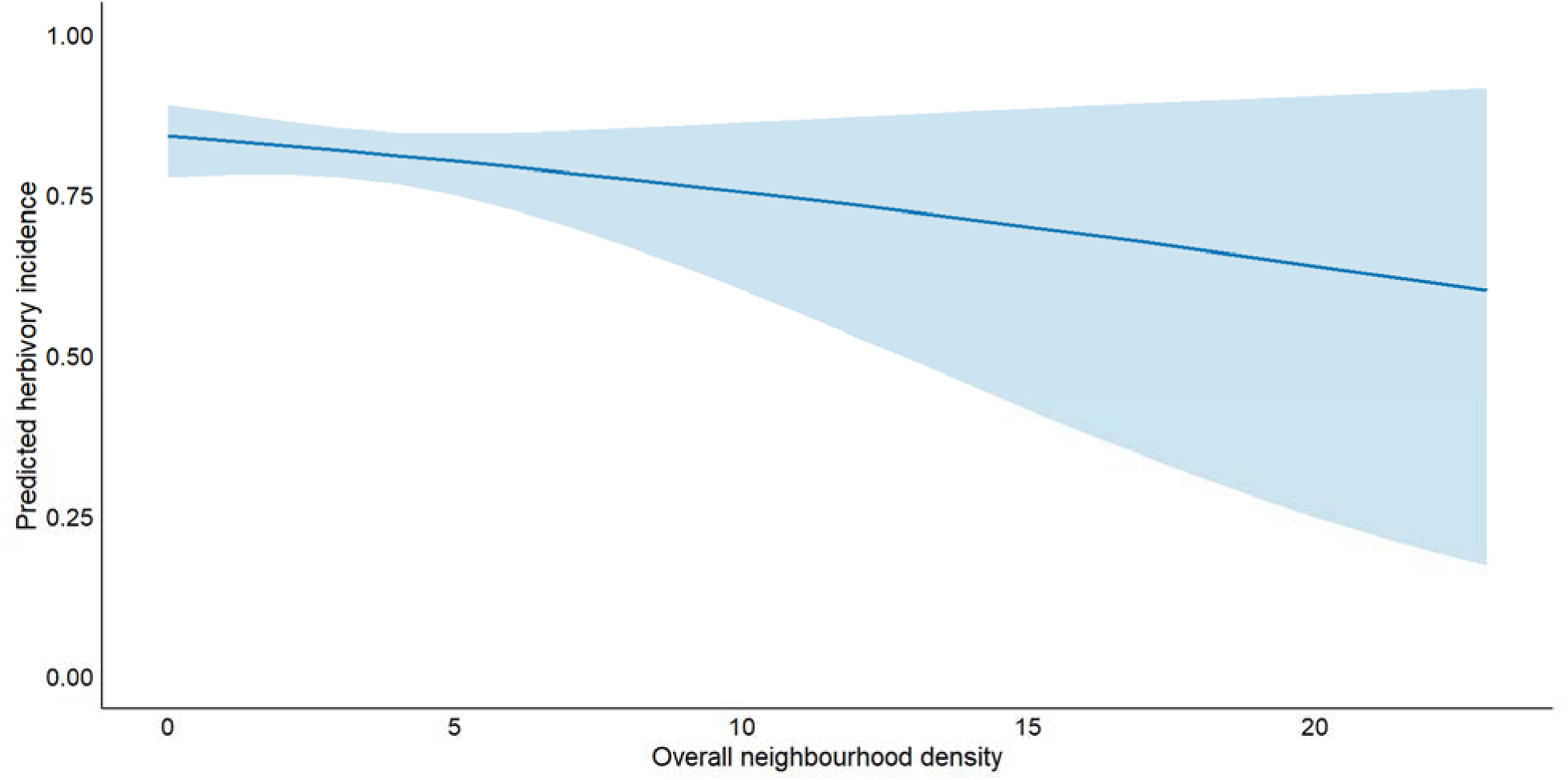
Predicted incidence of herbivory decreases with neighbourhood density. The shaded area shows 95% confidence intervals.

### 3.3 Species-specific responses to herbivory

Species identity, rather than coffee presence or absence, consistently emerged as the strongest and most consistent predictor of herbivory patterns. Both the incidence of herbivory and the extent of damage varied significantly among species (*P* < 0.001 in both cases; Figure 4; Table 1), regardless of treatment or neighbourhood density effects.

**Figure 4.**
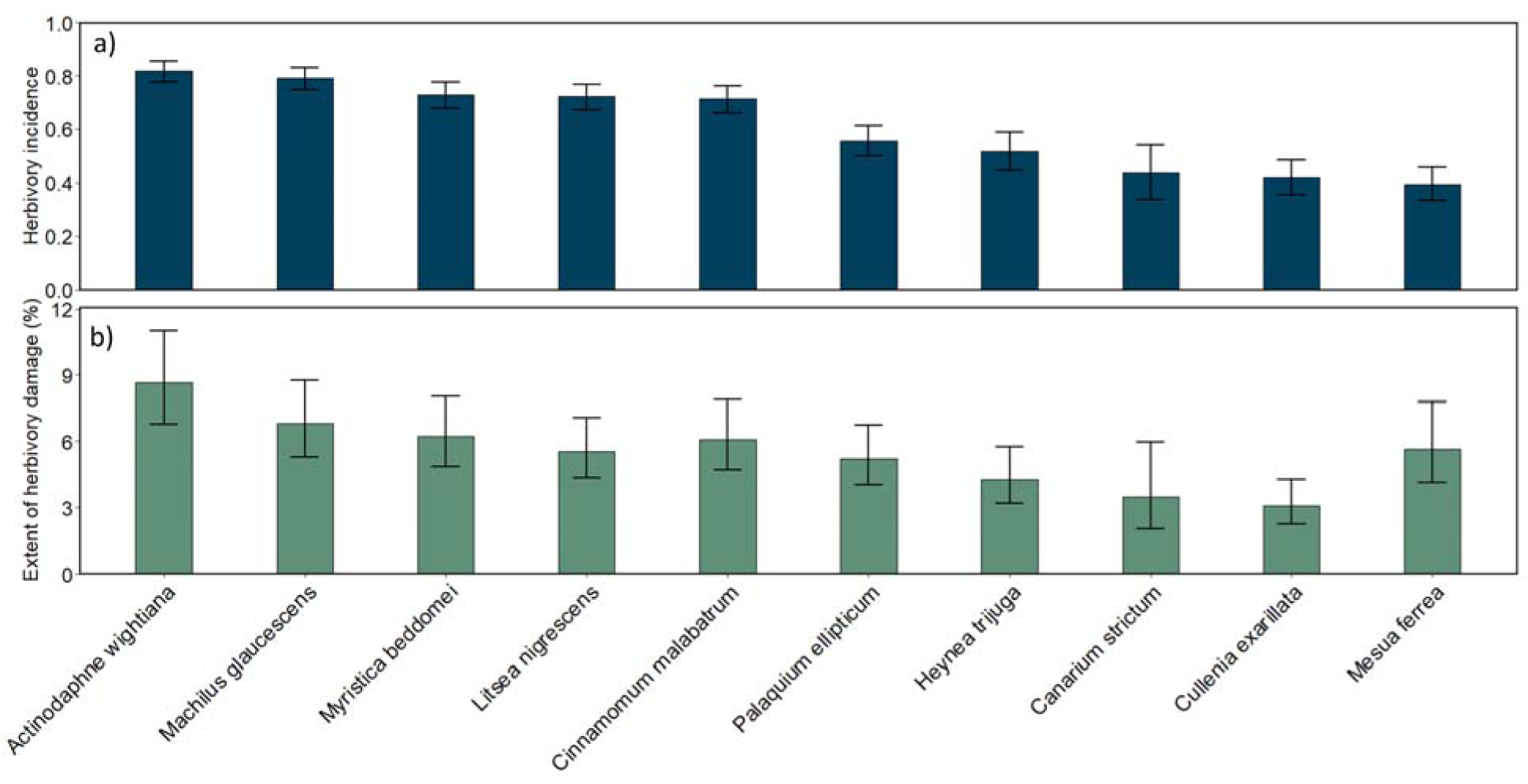
a) Predicted probability of herbivory incidence (% of leaves with any damage) across seedling species predicted from a GLMM with binomial distribution. b) Predicted extent of herbivory damage on seedlings of the different species, based on the linear mixed effect model. Bars show the model-predicted means for each species, averaged over treatment effects. Error bars represent 95% confidence intervals.

**Table 1.**
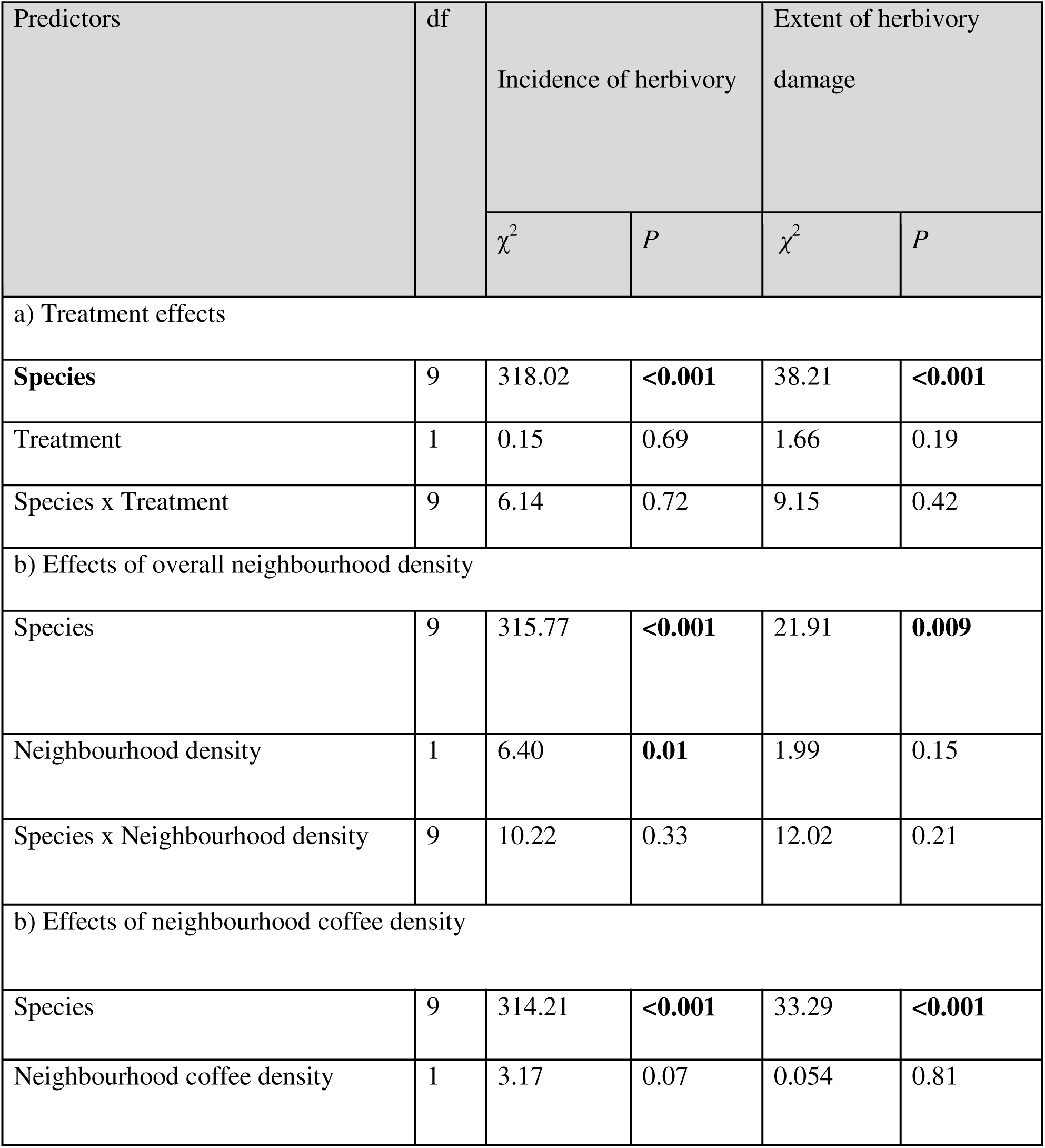
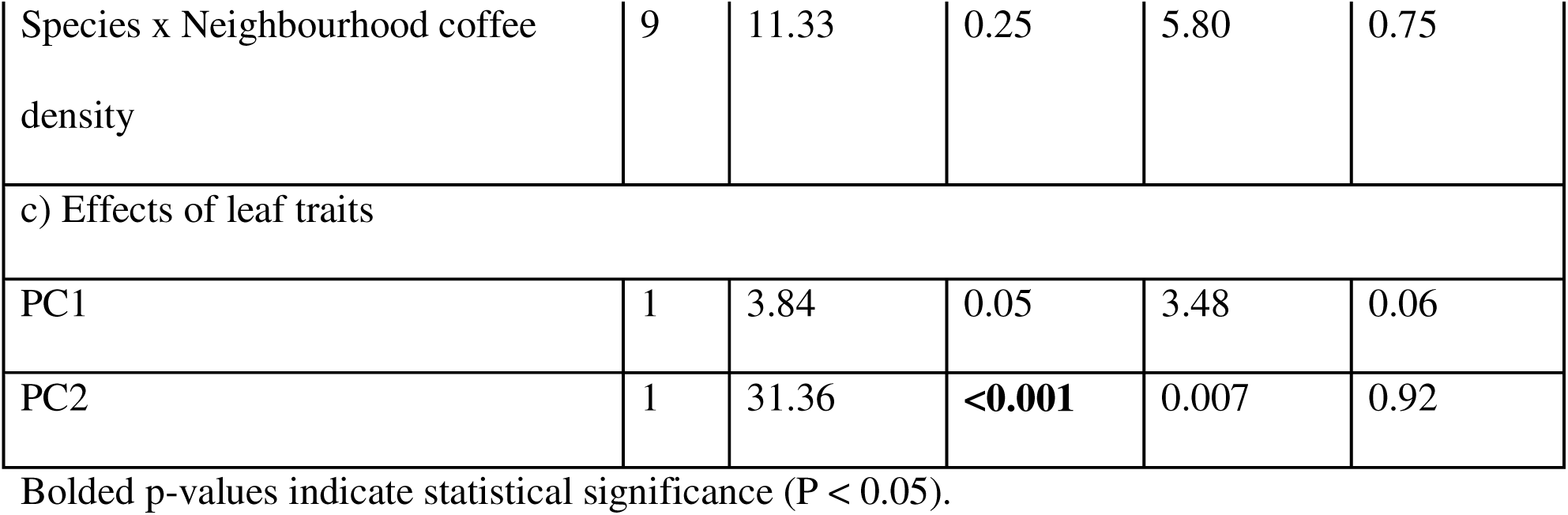
*ANOVA* summaries of the effects of treatments, neighbourhood density, and leaf traits on the incidence and extent of herbivory damage. The extent of damage was log-transformed for all analyses. Full model summaries can be found in the supplementary materials.

#### Role of plant traits

To test whether interspecific variation in herbivory could be explained by differences in leaf traits, we first examined the correlations among the six measured traits (Table S7). The correlation matrix revealed strong associations (Figure S4). Specific leaf area (SLA) was positively correlated with leaf nitrogen *(r = 0.64, df = 8)* and negatively correlated with leaf carbon-to-nitrogen ratio (r = - 0.88) and leaf dry matter content (LDMC; *r = – 0.83)*. Leaf nitrogen was negatively correlated with both LDMC *(r = –0.71)* and C: N ratio *(r = –0.90)*, while LDMC showed strong positive correlations with both leaf carbon *(r = 0.73)* and C: N ratio *(r = 0.83)*. The Principal Component Analysis (PCA) of leaf traits revealed two primary axes of trait variation that cumulatively explained over 80% of the total trait variation (PC1: 59.4%, PC2: 20.9%; Table S8; Figure S5). The first principal component (PC1) was strongly positively associated with leaf nitrogen (loading = 0.49) and SLA (0.49), and negatively correlated with leaf thickness (−0.41), LDMC (−0.38) and leaf C: N ratio, thus differentiating resource-acquisitive species (positive PC1 scores) from resource-conservative ones (low PC1 scores). PC2 (20.9% variance), on the other hand, was negatively correlated with leaf carbon (−0.85; Table S8), potentially reflecting leaf structural investment.

PC2 was strongly associated with herbivory incidence but not with the extent of herbivory damage. As PC2 increases (indicating lower leaf carbon content and structural investment), the incidence of herbivory increases χ² = 31.36, SE = 0.01; Figure 5b). PC1 showed a marginal negative association with herbivory incidence and extent of damage (incidence: χ² = 3.84, P = 0.05; damage: χ² = 3.48, P = 0.06; Figure 5a, Table S9 & S10). Counter to expectations, leaves with higher nitrogen content and specific leaf area (SLA), coupled with lower thickness, leaf dry matter content (LDMC), and C: N ratio, were associated with lower incidences of herbivory and extent of damage. However, the effect size was relatively small (Table 1).

**Figure 5.**
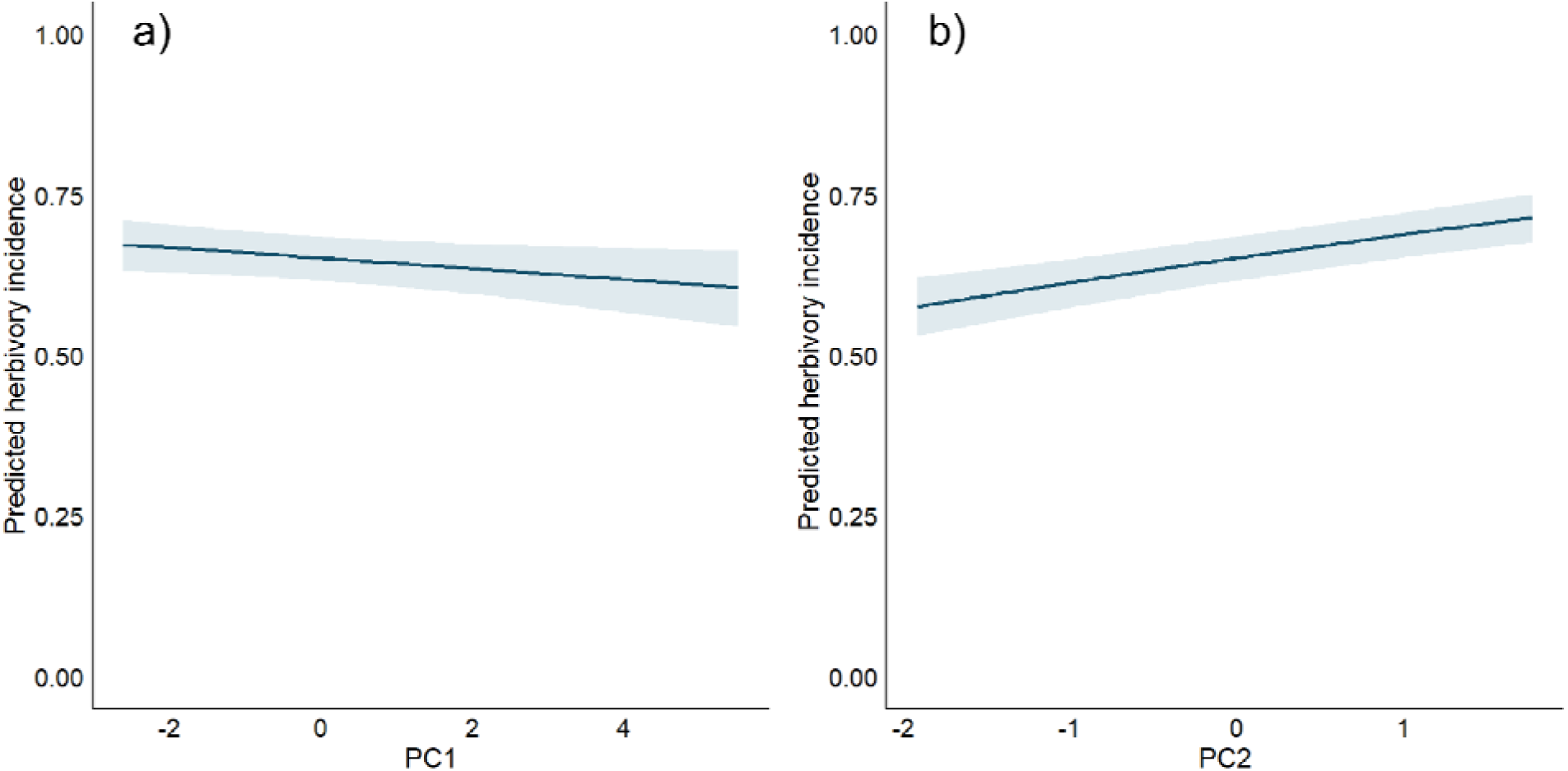
Predicted herbivory incidence from a generalised linear mixed model (binomial) plotted against (a) PC1 and (b) PC2 scores from a PCA of leaf traits. Shaded regions indicate 95% confidence intervals.

## 4. DISCUSSION

This study investigated the effects of removing the invasive understory species *Coffea canephora* on insect herbivory patterns in native tree seedlings in a rainforest restoration experiment. The removal of coffee had no detectable impact on herbivory patterns; however, increased neighbourhood densities of plants around focal individuals significantly reduced the incidence of herbivory. Another key finding was that species identity emerged as the primary driver of herbivory, with leaf carbon content playing a significant role in reducing the probability of herbivory incidence. Notably, none of the predictors influenced the extent of herbivory damage.

In support of neighbourhood effects shaping plant-insect interactions, as reported previously in the literature from both natural forests and experimental studies (Hambäck, Inouye, Andersson, & Underwood, 2014b; Otway, Hector, & Lawton, 2005b; Wenninger, Kim, Spiesman, & Gratton, 2016), we too found evidence for resource dilution effects in our restoration experiment, where higher local plant densities were associated with a decreased probability of herbivory incidence. However, once a seedling was attacked, the extent of herbivory damage was not related to neighbourhood density. This suggests that the initial decision by herbivores to attack a plant is influenced by the neighbourhood density, but the extent of feeding, once initiated, is determined by other factors. Our finding that the incidence and extent of herbivory damage can be driven by divergent mechanisms is consistent with Barker et al., (2022), where incidence was driven by general traits but not extent of damage, likely because herbivore consumption itself is influenced by other species-specific cues. Overall, these findings suggest that a deeper investigation of how physical and chemical cues and defences influence herbivore decisions and plant damage will yield further insights.

Despite the evidence of a dilution effect on the incidence of herbivory, the removal of coffee plants did not affect herbivory patterns. One potential explanation is that coffee itself does not appear to mediate such effects, likely because it does not differ strongly from other species in traits that herbivores cue into, such as structural characteristics or palatability. Further, the density of coffee plants in the local neighbourhood, by itself, had no detectable effect on either herbivory incidence or extent of damage, potentially indicating that the observed dilution effects arise from the combined presence of diverse neighbours rather than any unique trait of coffee. This aligns with the literature suggesting that the identity of neighbouring species is critical in determining the strength and direction of associational effects (Barbosa et al., 2009; Hambäck et al., 2014). Not all neighbours contribute equally; rather, the traits of the neighbour, their phylogenetic relatedness, and their palatability to shared herbivores can all play a role in mediating such effects (Moreira et al., 2017). In light of our findings, we encourage future studies to consider the traits of neighbouring plants too, as these have been previously shown to influence the behaviour and survival of focal plants (Barbosa et al., 2009).

Herbivory incidence and extent of damage varied significantly across our ten focal species. There is growing recognition that plant identity is often the dominant driver of herbivory in diverse tropical forests (Descombes, Kergunteuil, Glauser, Rasmann, & Pellissier, 2020; Loranger et al., 2012a; Pérez-Harguindeguy et al., 2003a). This underscores the fundamental role of evolutionary history and species-specific traits in driving plant-herbivore interactions (Endara, Forrister, & Coley, 2023). Significant variation in both incidence and extent of herbivore damage across species suggests that herbivore preferences or host specialisation are tightly linked to intrinsic plant traits rather than environmental modifications.

Previous studies have shown that the variation in physical traits and the elemental composition of leaves of different species can lead to different susceptibility to herbivores (Descombes et al., 2020; Loranger et al., 2012a; Pérez-Harguindeguy et al., 2003b). In our study, leaf physical and chemical trait-based analysis revealed that leaf carbon content significantly reduced the probability of herbivore attack but had no influence on the extent of herbivory damage among attacked plants. Species with low leaf carbon (a proxy for high nutritional value and low structural investment) were more likely to be attacked, while carbon-rich species were avoided. High leaf carbon content serves as a deterrent signal to herbivory (Shao, Cheng, Zhang, Xu, & Li, 2024), potentially indicating increased structural defences (lignin, cellulose) or reduced nutritional quality that herbivores can assess during initial plant selection (P. D. Coley, Bryant, & Chapin, 1985). This chemical signal appears to influence herbivore host selection decisions, with carbon-rich leaves being less likely to be chosen for consumption. Interestingly, while leaf carbon content influenced initial feeding decisions, it did not predict the extent of damage once attacked. The subsequent lack of relationship between these traits and the extent of damage suggests that defensive traits not measured in our study, such as secondary compounds or structural elements, may become more important determinants of feeding intensity once herbivores begin consuming leaf tissue. For example, Liu et al., (2023) show that compounds like hemicellulose, lignin, and tannins strongly influence feeding behaviour in evergreen species. In sum, early feeding decisions may be driven by coarse cues like leaf carbon (structural toughness), while sustained feeding depends on finer-scale drivers. Contrary to expectations and previous findings (Phyllis D. Coley, Bryant, & Chapin, 1985; Loranger et al., 2012b; Pérez-Harguindeguy et al., 2003a), we found that leaves with higher nitrogen content and specific leaf area (SLA), coupled with lower thickness, leaf dry matter content (LDMC), and C: N ratio, were linked to reduced herbivory incidence. One potential explanation is that high nitrogen content could be supporting the production of nitrogen-rich defensive compounds (Sun, Wang, Mur, Shen, & Guo, 2020). A deeper examination of how species’ defence traits vary with other leaf traits and jointly impact herbivore attacking and feeding behaviours could yield further insights. A caveat is that we quantified traits of adult trees of our target species and not from seedlings themselves, and it is likely that trait and herbivory relationships change based on plant ontogeny (Barton & Hanley, 2013; Barton & Koricheva, 2010).

In conclusion, our study explored how the removal of invasive coffee plants influenced insect herbivory on native tree seedlings in an ecological restoration experiment in the tropical forests of the Western Ghats. Removal of coffee plants did not affect either the incidence of herbivory or the extent of leaf damage. Thus, from a restoration perspective, the indirect effects of coffee on native seedlings are probably of far less consequence than its direct effects. At the same time, we found that a greater density of plants in the neighbourhood reduced the incidence of herbivory of focal plants. This sets up a potentially interesting trade-off from a restoration perspective: on one hand, plant neighbours might compete for resources with planted seedlings, negatively impacting their growth and survival. On the other hand, they might indirectly facilitate growth and survival by reducing herbivory through dilution effects. Targeted experiments can help shed light on such a tradeoff and its implications. Finally, herbivory varied substantially among species, and the interspecific differences in herbivory incidence were partly explained by species leaf traits. However, we only considered a small subset of functional traits and future research should consider examining other traits - in particular, understanding how herbivore impacts vary across early and late successional species may help inform restoration practices. While our study captures a snapshot of plant-insect herbivore interactions, long-term monitoring is required to understand how they change over time in restored systems.

## ACKNOWLEDGEMENTS

We thank Dr. Anand M. Osuri for facilitating access to the restoration experiment and for helping in developing the study design. We are also grateful to Dr. Sandeep Pulla for his guidance with statistical analyses. We sincerely acknowledge R. Rajesh, Srinivasan Kasinathan and other field staff from the Nature Conservation Foundation (NCF) for their dedicated support during fieldwork and data collection. We are grateful to all our colleagues for their suggestions and support during the study. We acknowledge the support provided to NCBS, TIFR by the Department of Atomic Energy, Government of India (Project Identification No. RTI 4006). Mahesh Sankaran acknowledges the support provided by the R. M. Tulpule Charitable Trust. Aparna Krishnan thanks Hindustan Unilever Ltd. for fellowship support. Mayank Kohli acknowledges the Department of Science & Technology, India, for fellowship support. Anand M Osuri, T R Shankar Raman and Divya Mudappa thank Rohini Nilekani Philanthropies and AMM Murugappa Chettiar Research Centre for funding and Parry Agro Industries Ltd for partnering in the rainforest restoration project.

## DATA AVAILABILITY STATEMENT

The data that support the findings of this study are available in the Dryad Digital Repository at *DOI: 10.5061/dryad.c59zw3rn2*. The data are currently under embargo and will be made publicly available upon article publication.

## AUTHOR CONTRIBUTIONS

LN, MK, MS, TRSR, and DM jointly conceived the research idea. LN, MK, MS, and AK contributed to the experimental setup and study design. LN and AM collected the field data. LN analysed the data and original draft of the manuscript, with MK, MS, TRSR, DM & AK providing critical feedback and making substantial contributions to the final version.

## CONFLICT OF INTEREST STATEMENT

The authors have no conflict of interest to declare.

## Notes

### Competing Interest Statement

The authors have declared no competing interest.

