## Supplementary Figures and Tables for "Insect herbivory on restored rainforest seedlings weakened by neighbours but unaffected by invasive coffee"

**Figure S1**. Experimental design set up. Both treatments (coffee present and coffee removed plots have 11 replicates each. Each plot was  20 × 20 m.

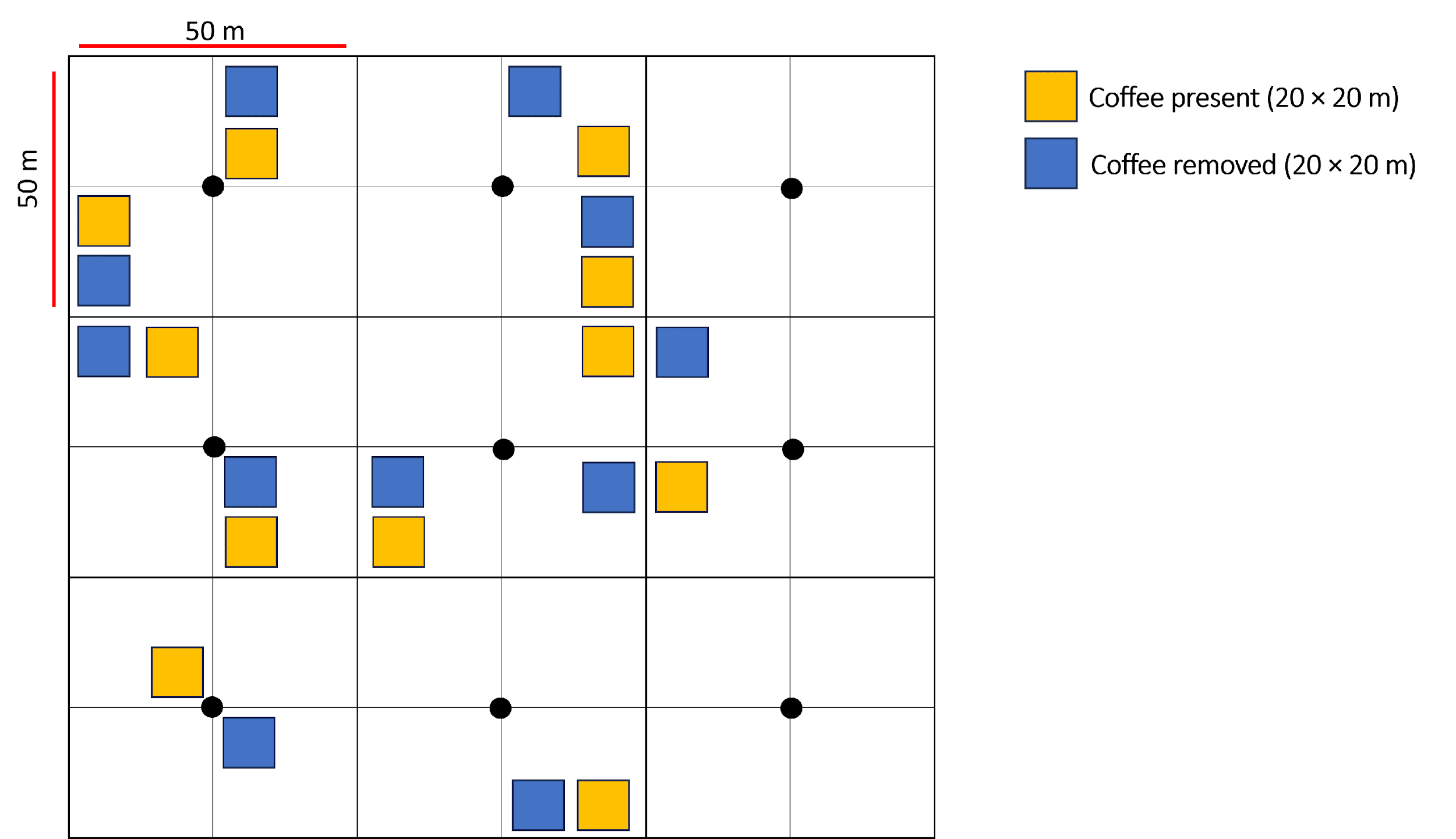

**Figure S2.** Distribution of the response variable (herbivory damage averaged across all the damaged leaves of an individual), pooled across all individuals (*N* = 824) and species in the study.

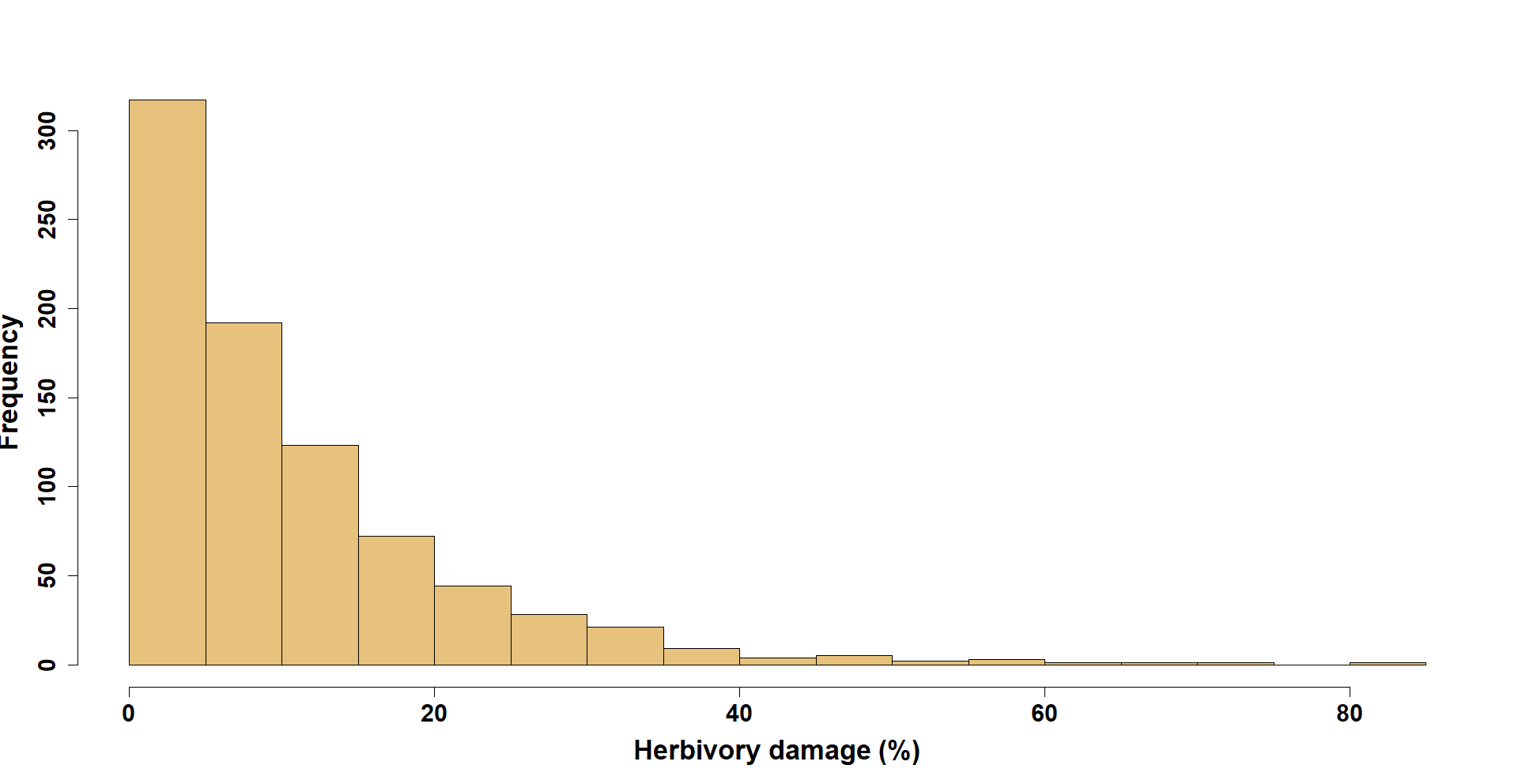

**Figure S3.** Distribution of the average neighbourhood density for two treatments (bars present mean ± SE).

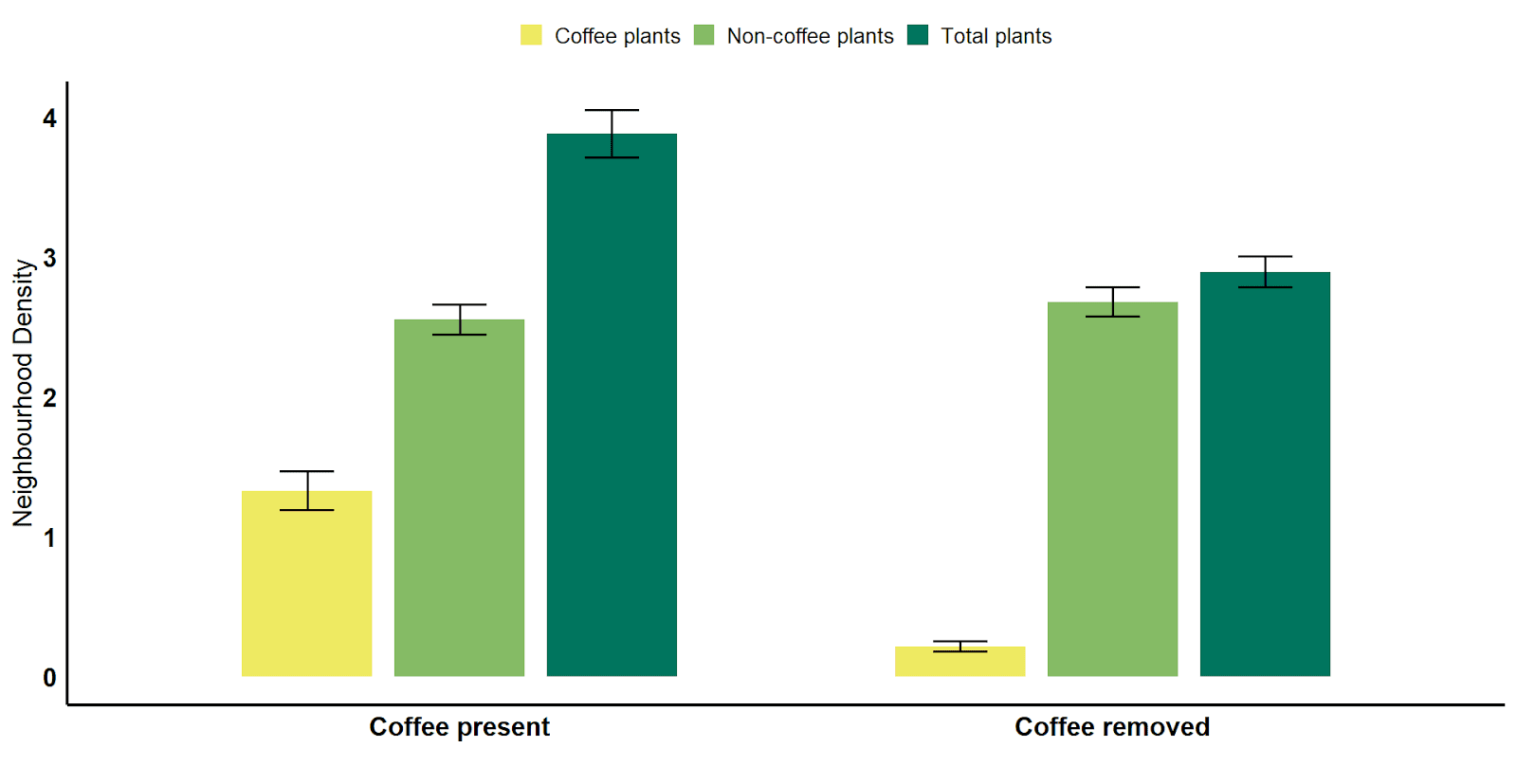

**Table S1.** Results of the generalised linear mixed model (binomial distribution) testing the effect of coffee removal on insect herbivory incidence on seedlings of directly-seeded plants.

**Model 1: Incidence	~ Treatment * Species + (1 | Plot /Individual )**

(Species abbreviations: ACT; *Actinodaphne wightiana*, CAN; *Canarium strictum*, CUL; *Cullenia exarillata*, HEY; *Heynea trijuga*, MES*; Mesua ferrea*, LIT; *Litsea nigrescens*,CIN; *Cinnamomum malabatrum*, MYR; *Myristica beddomei*, PAL*; Palaquium elipticum,* MAC; *Machilus glaucescens*)

|  | Estimate | Std. Error | z value | Pr(>\|z\|) |  |
| --- | --- | --- | --- | --- | --- |
| (Intercept) | 1.58671 | 0.1933 | 8.209 | 2.24E-16 | *** |
| Treatmentwithout_coffee | -0.1573 | 0.26925 | -0.584 | 0.559067 |  |
| Species_idCAN | -1.86214 | 0.35911 | -5.185 | 2.16E-07 | *** |
| Species_idCIN | -0.63471 | 0.22228 | -2.855 | 0.004298 | ** |
| Species_idCUL | -1.85489 | 0.22985 | -8.07 | 7.02E-16 | *** |
| Species_idHEY | -1.52922 | 0.22877 | -6.685 | 2.32E-11 | *** |
| Species_idLIT | -0.80121 | 0.21353 | -3.752 | 0.000175 | *** |
| Species_idMDAC | -0.53514 | 0.2235 | -2.394 | 0.016648 | * |
| Species_idMES | -2.09739 | 0.23406 | -8.961 | < 2e-16 | *** |
| Species_idPAL | -1.3936 | 0.20861 | -6.68 | 2.38E-11 | *** |
| Species_idMAC | -0.41182 | 0.2208 | -1.865 | 0.062158 | . |
| Treatmentwithout_coffee:Species_idCAN | 0.20509 | 0.47299 | 0.434 | 0.664583 |  |
| Treatmentwithout_coffee:Species_idCIN | 0.08043 | 0.31303 | 0.257 | 0.797237 |  |
| Treatmentwithout_coffee:Species_idCUL | 0.04018 | 0.33377 | 0.12 | 0.904185 |  |
| Treatmentwithout_coffee:Species_idHEY | 0.18973 | 0.34308 | 0.553 | 0.580245 |  |
| Treatmentwithout_coffee:Species_idLIT | 0.50117 | 0.30293 | 1.654 | 0.098046 | . |
| Treatmentwithout_coffee:Species_idMDAC | 0.03684 | 0.31693 | 0.116 | 0.907472 |  |
| Treatmentwithout_coffee:Species_idMES | 0.32553 | 0.32596 | 0.999 | 0.317952 |  |
| Treatmentwithout_coffee:Species_idPAL | 0.23348 | 0.29606 | 0.789 | 0.43033 |  |
| Treatmentwithout_coffee:Species_idMAC | 0.46988 | 0.32065 | 1.465 | 0.142811 |  |

Anova results of the model in Table S1 above

|  | Chisq | df | Pr(>Chisq) |  |
| --- | --- | --- | --- | --- |
| Treatment | 0.1553 | 1 | 0.6935 |  |
| Species_id | 318.0224 | 9 | <2e-16 | *** |
| Treatment: Species_id | 6.1475 | 9 | 0.7251 |  |

**Table S2.** Results of the linear mixed model (normal distribution) testing the effect of coffee removal on insect herbivory damage on seedlings of directly-seeded plants. Herbivory data has been log-transformed.

**Model 2: Herbivory damage ~ Treatment * Species + (1|Plot)**

|  | Estimate | Std. Error | t value |
| --- | --- | --- | --- |
| (Intercept) | 2.06347 | 0.17436 | 11.834 |
| Treatmentwithout_coffee | 0.18512 | 0.24659 | 0.751 |
| Species_idCAN | -0.97829 | 0.4393 | -2.227 |
| Species_idCIN | -0.32312 | 0.25427 | -1.271 |
| Species_idCUL | -1.23644 | 0.26999 | -4.58 |
| Species_idHEY | -0.51147 | 0.26594 | -1.923 |
| Species_idLIT | -0.45981 | 0.24643 | -1.866 |
| Species_idMDAC | -0.49846 | 0.24642 | -2.023 |
| Species_idMES | -0.29991 | 0.29037 | -1.033 |
| Species_idPAL | -0.31923 | 0.25014 | -1.276 |
| Species_idMAC | -0.05892 | 0.24643 | -0.239 |
| Treatmentwithout_coffee:Species_idCAN | 0.14042 | 0.59904 | 0.234 |
| Treatmentwithout_coffee:Species_idCIN | -0.05458 | 0.35963 | -0.152 |
| Treatmentwithout_coffee:Species_idCUL | 0.41352 | 0.40991 | 1.009 |
| Treatmentwithout_coffee:Species_idHEY | -0.38081 | 0.38551 | -0.988 |
| Treatmentwithout_coffee:Species_idLIT | 0.03135 | 0.3485 | 0.09 |
| Treatmentwithout_coffee:Species_idMDAC | 0.34611 | 0.35515 | 0.975 |
| Treatmentwithout_coffee:Species_idMES | -0.24307 | 0.40645 | -0.598 |
| Treatmentwithout_coffee:Species_idPAL | -0.36977 | 0.35777 | -1.034 |
| Treatmentwithout_coffee:Species_idMAC | -0.35723 | 0.35411 | -1.009 |

ANOVA results of the model in Table S2 above.

|  | Chisq | df | Pr(>Chisq) |  |
| --- | --- | --- | --- | --- |
| (Intercept) | 140.0527 | 1 | < 2.2e-16 | *** |
| Treatment | 0.5636 | 1 | 0.452821 |  |
| Species_id | 28.8213 | 9 | 0.000695 | *** |
| Treatment: Species_id | 9.1599 | 9 | 0.422645 |  |

**Table S3.** Results of the generalised linear mixed model (binomial distribution) testing the effect of overall neighbourhood density on insect herbivory incidence on seedlings of directly-seeded plants. Plot and individual are added as nested random effects.

**Model 3: Incidence ~ Overall neighbourhood density * Species + (1 | Plot /Individual)**

|  | Estimate | Std. Error | z value | Pr(>\|z\|) |  |
| --- | --- | --- | --- | --- | --- |
| (Intercept) | 1.6802 | 0.21591 | 7.782 | 7.13E-15 | *** |
| Species_idCAN | -1.95602 | 0.36195 | -5.404 | 6.51E-08 | *** |
| Species_idCIN | -0.76749 | 0.25762 | -2.979 | 0.00289 | ** |
| Species_idCUL | -1.76833 | 0.28337 | -6.24 | 4.36E-10 | *** |
| Species_idHEY | -1.1564 | 0.28189 | -4.102 | 4.09E-05 | *** |
| Species_idLIT | -0.49426 | 0.25251 | -1.957 | 0.0503 | . |
| Species_idMDAC | -0.59723 | 0.26064 | -2.291 | 0.02194 | * |
| Species_idMES | -1.94652 | 0.27519 | -7.073 | 1.51E-12 | *** |
| Species_idPAL | -1.44678 | 0.2587 | -5.592 | 2.24E-08 | *** |
| Species_idMAC | -0.26256 | 0.27012 | -0.972 | 0.33106 |  |
| total_neighbourhood_density | -0.05527 | 0.0513 | -1.078 | 0.28123 |  |
| Species_idCAN:total_neighbourhood_density | 0.05783 | 0.07118 | 0.813 | 0.41649 |  |
| Species_idCIN:total_neighbourhood_density | 0.05763 | 0.06065 | 0.95 | 0.34194 |  |
| Species_idCUL:total_neighbourhood_density | -0.01667 | 0.06977 | -0.239 | 0.81112 |  |
| Species_idHEY:total_neighbourhood_density | -0.08408 | 0.06873 | -1.223 | 0.22118 |  |
| Species_idLIT:total_neighbourhood_density | -0.02179 | 0.06074 | -0.359 | 0.71974 |  |
| Species_idMDAC:total_neighbourhood_density | 0.03217 | 0.05754 | 0.559 | 0.57613 |  |
| Species_idMES:total_neighbourhood_density | 0.01218 | 0.06405 | 0.19 | 0.84917 |  |
| Species_idPAL:total_neighbourhood_density | 0.05415 | 0.05937 | 0.912 | 0.3617 |  |
| Species_idMAC:total_neighbourhood_density | 0.02579 | 0.06182 | 0.417 | 0.67658 |  |

Anova results of the model in Table S3 above

|  | Chisq | df | Pr(>Chis) |  |
| --- | --- | --- | --- | --- |
| Species_id | 315.832 | 9 | < 2e-16 | *** |
| total_neighbourhood_density | 6.4027 | 1 | 0.01139 | * |
| Species_id:total_neighbourhood_density | 10.2235 | 9 | 0.33269 |  |

**Table S4.** Results of the linear mixed model (normal distribution) testing the effect of overall neighbourhood density on insect herbivory damage on seedlings of directly-seeded plants. Herbivory data has been log-transformed.

**Model 4: Herbivory damage ~ Overall neighbourhood density * Species + (1|Plot)**

|  | Estimate | Std. Error | t value |
| --- | --- | --- | --- |
| (Intercept) | 1.90977 | 0.21399 | 8.925 |
| Species_idCAN | -0.60136 | 0.437567 | -1.374 |
| Species_idCIN | -0.18402 | 0.284964 | -0.646 |
| Species_idCUL | -1.11556 | 0.332916 | -3.351 |
| Species_idHEY | -0.43156 | 0.314162 | -1.374 |
| Species_idLIT | -0.25204 | 0.279626 | -0.901 |
| Species_idMDAC | 0.182632 | 0.279051 | 0.654 |
| Species_idMES | 0.082871 | 0.333543 | 0.248 |
| Species_idPAL | -0.33235 | 0.307028 | -1.082 |
| Species_idMAC | -0.14206 | 0.292119 | -0.486 |
| total_neighbourhood_density | 0.076619 | 0.054277 | 1.412 |
| Species_idCAN:total_neighbourhood_density | -0.0866 | 0.085651 | -1.011 |
| Species_idCIN:total_neighbourhood_density | -0.05236 | 0.068126 | -0.769 |
| Species_idCUL:total_neighbourhood_density | 0.009788 | 0.083992 | 0.117 |
| Species_idHEY:total_neighbourhood_density | -0.08197 | 0.085735 | -0.956 |
| Species_idLIT:total_neighbourhood_density | -0.0582 | 0.070357 | -0.827 |
| Species_idMDAC:total_neighbourhood_density | -0.15098 | 0.063845 | -2.365 |
| Species_idMES:total_neighbourhood_density | -0.14638 | 0.075931 | -1.928 |
| Species_idPAL:total_neighbourhood_density | -0.05737 | 0.070063 | -0.819 |
| Species_idMAC:total_neighbourhood_density | -0.03302 | 0.069334 | -0.476 |

ANOVA results of the model in Table S4 above.

|  | Chisq | df | Pr(>Chisq) |  |
| --- | --- | --- | --- | --- |
| (Intercept) | 79.6484 | 1 | < 2.2e-16 | *** |
| Species_id | 21.9153 | 9 | 0.009151 | ** |
| total_neighbourhood_density | 1.9927 | 1 | 0.15806 |  |
| Species_id:total_neighbourhood_density | 12.027 | 9 | 0.211789 |  |

**Table S5.** Results of the generalised linear mixed model (binomial distribution) testing the effect of neighbourhood coffee density on insect herbivory incidence on seedlings of directly-seeded plants. Plot and individual are added as nested random effects.

**Model 5: Incidence ~ Neighbourhood coffee density * Species + (1 | Plot /Individual)**

|  | Estimate | Std. Error | z value | Pr(>\|z\|) |  |
| --- | --- | --- | --- | --- | --- |
| (Intercept) | 1.46369 | 0.14408 | 10.159 | < 2e-16 | *** |
| Species_idCAN | -1.77058 | 0.24989 | -7.085 | 1.39E-12 | *** |
| Species_idCIN | -0.4982 | 0.16952 | -2.939 | 0.00329 | ** |
| Species_idCUL | -1.75355 | 0.18014 | -9.735 | < 2e-16 | *** |
| Species_idHEY | -1.20119 | 0.18867 | -6.367 | 1.93E-10 | *** |
| Species_idLIT | -0.4561 | 0.16247 | -2.807 | 0.00499 | ** |
| Species_idMDAC | -0.48931 | 0.17134 | -2.856 | 0.00429 | ** |
| Species_idMES | -1.8227 | 0.17969 | -10.143 | < 2e-16 | *** |
| Species_idPAL | -1.21934 | 0.16145 | -7.552 | 4.28E-14 | *** |
| Species_idMAC | -0.11169 | 0.17265 | -0.647 | 0.51771 |  |
| Neighbourhood_coffee_density | 0.08401 | 0.11035 | 0.761 | 0.44645 |  |
| Species_idCAN:Neighbourhood_coffee_density | -0.05138 | 0.12136 | -0.423 | 0.67201 |  |
| Species_idCIN:Neighbourhood_coffee_density | -0.13391 | 0.11592 | -1.155 | 0.24801 |  |
| Species_idCUL:Neighbourhood_coffee_density | -0.12941 | 0.15709 | -0.824 | 0.41005 |  |
| Species_idHEY:Neighbourhood_coffee_density | -0.26023 | 0.12801 | -2.033 | 0.04206 | * |
| Species_idLIT:Neighbourhood_coffee_density | -0.18725 | 0.12158 | -1.54 | 0.12352 |  |
| Species_idMDAC:Neighbourhood_coffee_density | -0.06667 | 0.116 | -0.575 | 0.56548 |  |
| Species_idMES:Neighbourhood_coffee_density | -0.14124 | 0.1222 | -1.156 | 0.24778 |  |
| Species_idPAL:Neighbourhood_coffee_density | -0.0982 | 0.11602 | -0.846 | 0.39732 |  |
| Species_idMAC:Neighbourhood_coffee_density | -0.1226 | 0.11777 | -1.041 | 0.29787 |  |

Anova results of the model in Table S5 above

|  | Chisq | df | Pr(>Chis) |  |
| --- | --- | --- | --- | --- |
| Species_id | 314.2186 | 9 | < 2e-16 | *** |
| Neighbourhood_coffee_density | 3.1796 | 1 | 0.07456 |  |
| Species_id:Neighbourhood_coffee_density | 11.3325 | 9 | 0.2536 |  |

**Table S6.** Results of the linear mixed model (normal distribution) testing the effect of neighbourhood coffee density on insect herbivory damage on seedlings of directly-seeded plants. Herbivory data has been log-transformed.

**Model 6: Herbivory damage ~ Neighbourhood coffee density * Species + (1|Plot)**

|  | Estimate | Std. Error | t value |
| --- | --- | --- | --- |
| (Intercept) | 2.168893 | 0.13615 | 15.93 |
| Species_idCAN | -0.9077 | 0.318342 | -2.851 |
| Species_idCIN | -0.41242 | 0.193873 | -2.127 |
| Species_idCUL | -1.10657 | 0.220078 | -5.028 |
| Species_idHEY | -0.71887 | 0.214375 | -3.353 |
| Species_idLIT | -0.40527 | 0.18677 | -2.17 |
| Species_idMDAC | -0.3361 | 0.192091 | -1.75 |
| Species_idMES | -0.36226 | 0.223259 | -1.623 |
| Species_idPAL | -0.54238 | 0.195966 | -2.768 |
| Species_idMAC | -0.29377 | 0.192048 | -1.53 |
| Neighbourhood_coffee_density | -0.02598 | 0.111118 | -0.234 |
| Species_idCAN:Neighbourhood_coffee_density | 0.030098 | 0.132594 | 0.227 |
| Species_idCIN:Neighbourhood_coffee_density | 0.082698 | 0.122105 | 0.677 |
| Species_idCUL:Neighbourhood_coffee_density | 0.028282 | 0.189445 | 0.149 |
| Species_idHEY:Neighbourhood_coffee_density | 0.047635 | 0.160985 | 0.296 |
| Species_idLIT:Neighbourhood_coffee_density | -0.07492 | 0.131928 | -0.568 |
| Species_idMDAC:Neighbourhood_coffee_density | 0.006708 | 0.119267 | 0.056 |
| Species_idMES:Neighbourhood_coffee_density | -0.0447 | 0.130238 | -0.343 |
| Species_idPAL:Neighbourhood_coffee_density | 0.05406 | 0.124008 | 0.436 |
| Species_idMAC:Neighbourhood_coffee_density | 0.080911 | 0.124036 | 0.652 |

ANOVA results of the model in Table S6 above.

|  | Chisq | df | Pr(>Chisq) |  |
| --- | --- | --- | --- | --- |
| (Intercept) | 253.7687 | 1 | < 2.2e-16 | *** |
| Species_id | 33.2946 | 9 | 0.000119 | *** |
| Neighbourhood_coffee_density | 0.0547 | 1 | 0.815151 |  |
| Species_id:Neighbourhood_coffee_density | 5.8024 | 9 | 0.759525 |  |

**Table S7.** Average trait values for the ten directed seedlings.

| Species | Leaf thickness | Leaf dry matter content (LDMC) | Specific leaf area (SLA) | leaf N (%) | leafc (%) | leafcn(%) |
| --- | --- | --- | --- | --- | --- | --- |
| *Actinodaphne wightiana* | 0.64 | 0.46 | 106.85 | 1.33 | 46.21 | 34.51 |
| *Canarium strictum* | 0.39 | 0.34 | 110.22 | 1.96 | 45.62 | 23.26 |
| *Cinnamomum malabatrum* | 0.61 | 0.62 | 86.79 | 1.54 | 46.62 | 30.27 |
| *Cullenia exarillata* | 0.60 | 0.45 | 87.33 | 1.22 | 46.48 | 38.06 |
| *Heynea trijuga* | 0.28 | 0.26 | 214.73 | 2.64 | 47.28 | 17.90 |
| *Litsea nigrescens* | 0.67 | 0.39 | 79.97 | 1.61 | 45.43 | 28.21 |
| *Myristica beddomei* | 0.57 | 0.36 | 102.45 | 1.27 | 48.11 | 37.88 |
| *Mesua ferrea* | 0.80 | 0.48 | 86.56 | 1.06 | 47.82 | 44.85 |
| *Palaquium elipticum* | 0.25 | 0.40 | 137.71 | 1.35 | 46.36 | 34.34 |
| *Machilus glaucescens* | 0.75 | 0.38 | 105.87 | 1.44 | 46.89 | 32.56 |

**Figure S4**. Correlation matrix among leaf trait variables

(Species abbreviations: ldmc; *Leaf dry matter content*, avg_thickness; *Leaf thickness*, sla; *Specific leaf area*, leafc; *Leaf carbon*, leafn*; Leaf nitrogen*, leafcn; *Leaf C:N*)

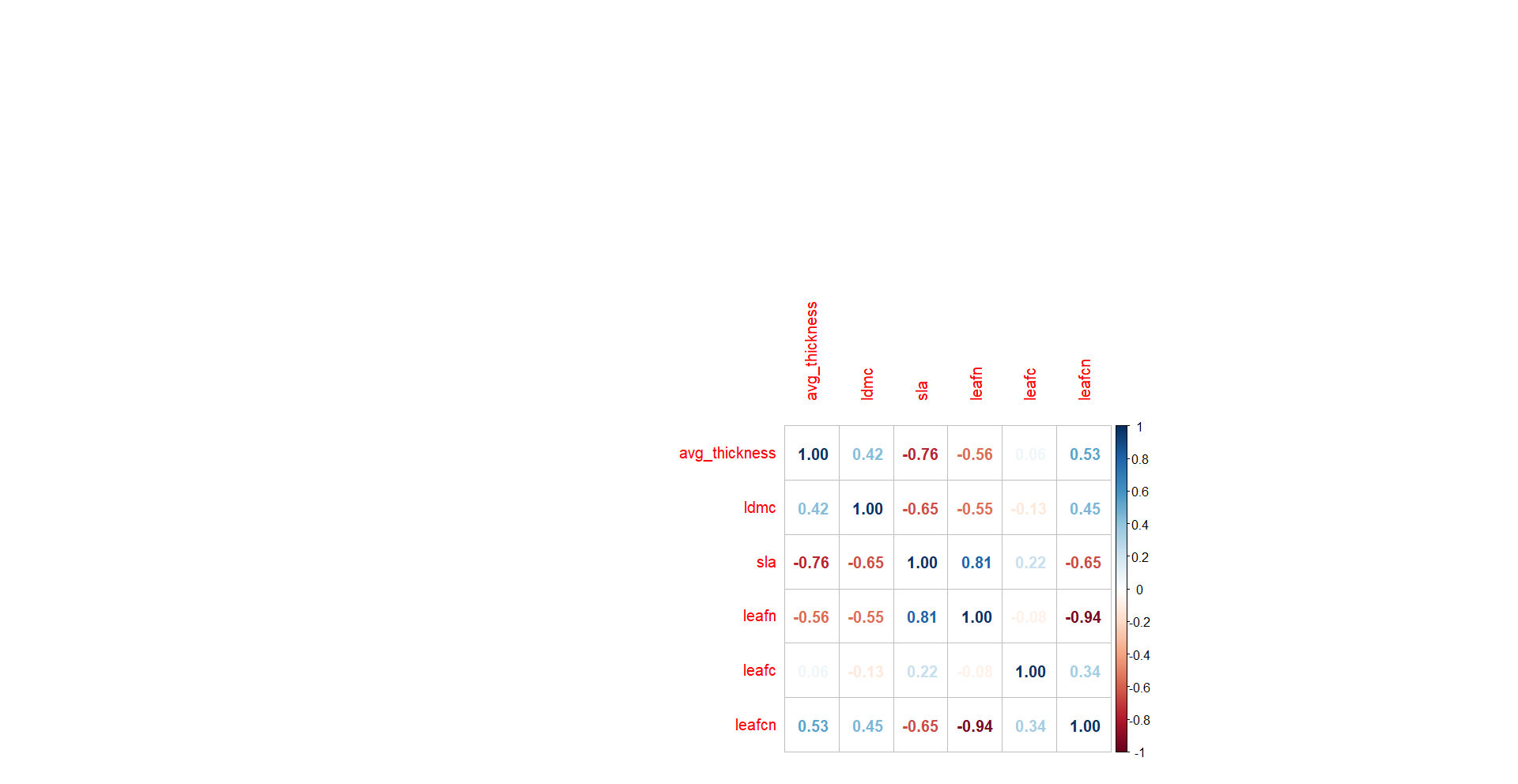

**Table S8**. Results of principal components (PC) analysis of leaf trait variables in the ten study species.

|  | **PC1** | **PC2** | **PC3** | **PC4** | **PC5** | **PC6** |
| --- | --- | --- | --- | --- | --- | --- |
| ‍**Variables** |  |  |  |  |  |  |
| Specific leaf area | 0.485963 | -0.2568 | 0.174375 | 0.084859 | -0.78193 | -0.22108 |
| Leaf carbon | -0.02429 | -0.8528 | -0.03398 | 0.387358 | 0.330259 | -0.10899 |
| Leaf nitrogen | 0.492828 | 0.108355 | -0.22913 | 0.403805 | 0.058288 | 0.725551 |
| Leaf thickness | -0.40948 | 0.029488 | -0.77691 | 0.271032 | -0.38208 | -0.09176 |
| Leaf dry matter content | -0.37675 | 0.26366 | 0.518916 | 0.709335 | -0.12686 | -0.00418 |
| Leaf C:N | -0.45908 | -0.35308 | 0.207533 | -0.32106 | -0.33771 | 0.635911 |
| ‍**Estimates** |  |  |  |  |  |  |
| Standard deviation | 1.8883 | 1.1208 | 0.7637 | 0.70223 | 0.29463 | 0.1226 |
| Proportion of Variance | 0.5943 | 0.2094 | 0.0972 | 0.08219 | 0.01447 | 0.0025 |
| Cumulative Proportion | 0.5943 | 0.8036 | 0.9008 | 0.98303 | 0.9975 | 1 |

**Figure S5.** Principal component analysis

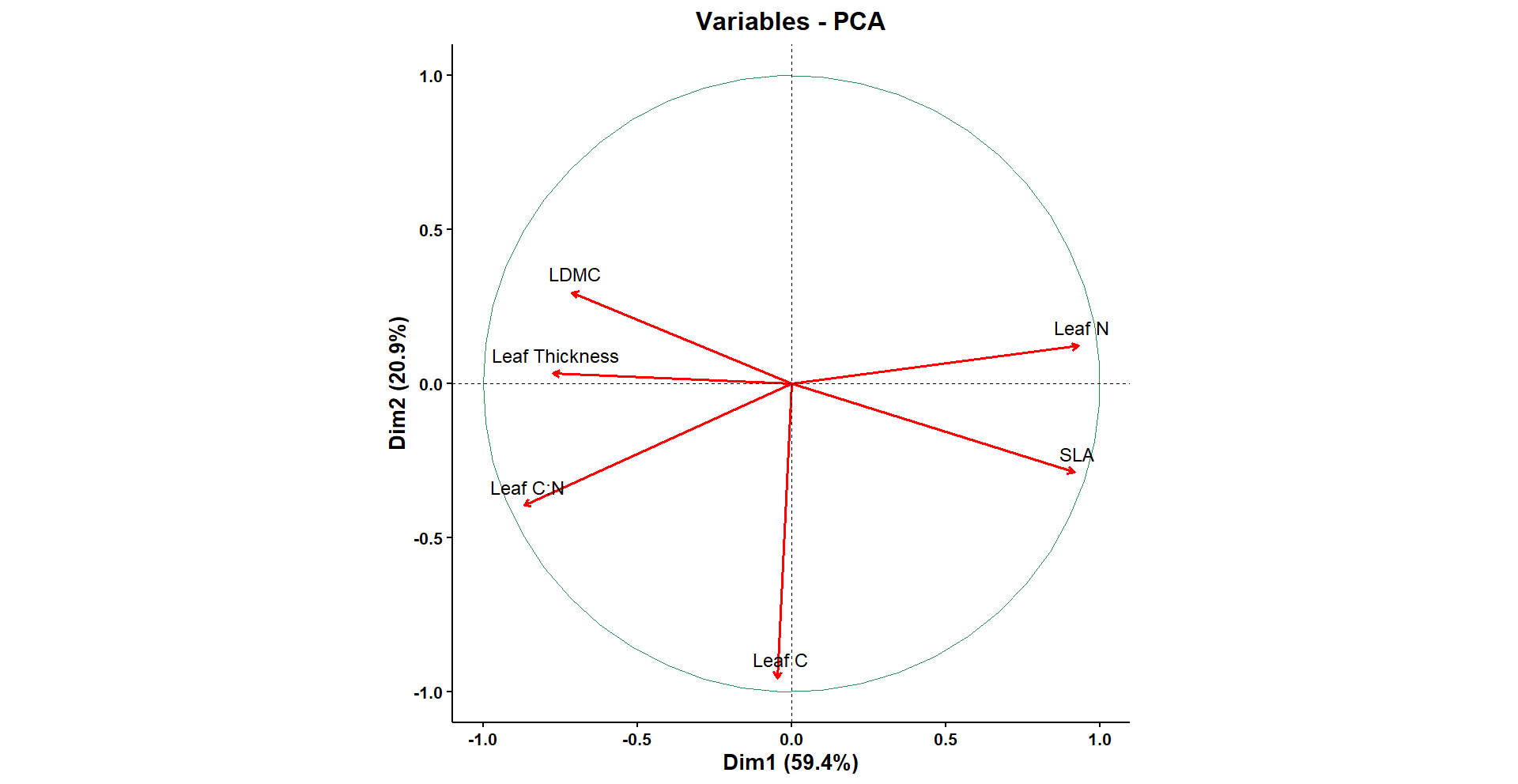

**Table S9.** Results of the generalised linear mixed model testing the effect of measured leaf traits on insect herbivory incidence on seedlings of directly seeded plants.

**Model 7: Incidence	~ (PC1+PC2) + (1 | Plot /Individual)**

|  | Estimate | Std. Error | z value | Pr(>\|z\|) |  |
| --- | --- | --- | --- | --- | --- |
| (Intercept) | 0.62702 | 0.07729 | 8.112 | 4.97E-16 | *** |
| PC1 | -0.03595 | 0.01834 | -1.96 | 0.05 | . |
| PC2 | 0.1657 | 0.02959 | 5.6 | 2.14E-08 | *** |

Anova results of the model in Table S8 above

|  | Chisq | df | Pr(>Chisq) |  |
| --- | --- | --- | --- | --- |
| PC1 | 3.8402 | 1 | 0.05004 | . |
| PC2 | 31.363 | 1 | 2.14E-08 | *** |

**Table S10.** Results of the linear mixed model (normal distribution) testing the effect of plant traits on insect herbivory damage on seedlings of directly-seeded plants. Herbivory data has been log-transformed.

**Model 8: Herbivory damage ~ (PC1+PC2) + (1|Plot)**

|  | Estimate | Std. Error | t value |
| --- | --- | --- | --- |
| (Intercept) | 1.730023 | 0.047146 | 36.695 |
| PC1 | -0.04567 | 0.02447 | -1.866 |
| PC2 | 0.003527 | 0.039699 | 0.089 |

ANOVA results of the model in Table S9 above.

|  | Chisq df Pr(>Chisq) | |
| --- | --- | --- |
| (Intercept) | 1346.5491 1 < 2e-16 | *** |
| PC1 | 3.4834 1 0.06199 |  |
| PC2 | 0.0079 1 0.92920 | |
